# Platelet response to influenza vaccination reflects effects of aging

**DOI:** 10.1101/2022.04.06.487196

**Authors:** A. Konstorum, S. Mohanty, Y. Zhao, A. Melillo, B. Vander Wyk, A. Nelson, S. Tsang, T.P. Blevins, R.B. Belshe, D.G. Chawla, M.T. Rondina, R.R. Montgomery, H.G. Allore, S.H. Kleinstein, A.C. Shaw

## Abstract

Platelets are uniquely positioned as mediators of not only hemostasis but also innate immunity, but how age and alterations in functional status such as frailty influence platelet function during an immune response remains unclear. We assessed the platelet transcriptome in younger (age 21-35) and older (age ≥ 65) adults (including frail and non-frail individuals) following influenza vaccination. Prior to vaccination, we identified an age- and frailty-associated increase in expression of platelet activation and mitochondrial RNAs and decrease in RNAs encoding proteins mediating translation. Using tensor decomposition analysis, we also elucidated dynamic post-vaccination platelet activation and translation signatures associated with age and frailty. At the protein level, enhanced platelet activation was found in non-frail older adults, compared to young individuals both prior to and post-vaccine; but frail adults showed decreased platelet activation compared to non-frail that could reflect the influence of decreased translation RNA expression. Our results reveal an age-dependent alteration in platelet function prior to and post-vaccination that may contribute to age-associated chronic inflammation.

---

Substantial evidence indicates that aging is associated with a heightened systemic chronic inflammatory state^1,2^; the consequences of such dysregulated inflammation include heightened risks for age-related diseases such as metabolic syndromes, diabetes, cardiovascular disease, and neurodegenerative disease^3,4^. The mechanisms underlying the development of age-related chronic inflammation remain incompletely understood, but are manifested by elevated levels of acute phase reactants, clotting factors, and cytokines. Several interrelated mechanisms underlying this inflammation have been hypothesized, including activation of innate immune pattern recognition receptor (PRR) signaling by age-related increases in damage-associated molecular patterns (DAMPs)^5^, the senescence-associated secretion phenotype (SASP), a secretome including pro-inflammatory cytokines induced by cellular damage^6^, and altered innate immune PRR function^7–10^.

Platelets are uniquely positioned as mediators of the age-associated chronic inflammatory milieu and are principally known as mediators of coagulation and not a typical focus of studies on the biology of aging. These anucleate cell fragments develop in the bone marrow and may reflect systemic inflammation in circulation. Several studies show that platelet counts decrease with age^11,12^, and that platelets from older individuals show elevated levels of the pro-thrombotic platelet factor 4 and β-thromboglobulin. In addition, responsiveness to *in vitro* stimuli promoting platelet aggregation (such as ADP) has also been reported to increase with age^13–17^. Notably, platelets secrete cytokines and other inflammatory mediators and express a full complement of innate immune PRRs including Toll-like Receptors (TLRs)^18,19^. Moreover, platelets contain mRNA, mitochondria and mitochondrial DNA, and machinery for post-transcriptional regulation of RNA expression. Our previous studies of PBMC transcriptomics to elucidate the effects of age on gene expression signatures of influenza vaccine response revealed modulation of platelet activation pathways, suggesting a role for platelet genes and the presence of platelet-leukocyte aggregates^20^. We therefore undertook the present study with a focus on alterations in the platelet transcriptome in response to influenza vaccination, coupled with assessment of platelet activation status at the protein level in cohorts of young and older adults.

## Results

We enrolled 28 young adults (age 21-35 years), 20 community-dwelling older adults (age ≥ 65 years) (Older (Comm)), and 17 older adults (age ≥ 65 years) (Older (SNF)) who were residents of a skilled nursing facility (SNF) in greater New Haven, Connecticut. As expected, Older (Comm) adults had more comorbid medical conditions and used more medications than young individuals, and the Older (SNF) group showed a further increase in comorbidities and numbers of medications (Table 1). The functional status of the Older (Comm) and Older (SNF) adults was assessed using an operational definition of frailty, a geriatric syndrome of decreased reserve in response to physiologic stress that is associated with adverse healthcare outcomes, increased disability, and mortality^21^. This frailty assessment used a five-point scale that included measurements of grip strength and gait speed, as well as assessments of unintentional weight loss, decreased physical activity, and exhaustion using validated instruments; individuals meeting criteria for three or more of these criteria are considered frail, 1 or 2 criteria pre-frail, and zero non-frail^21^. The Older (Comm) individuals were non-frail except for four individuals who were classified as pre-frail, while the Older (SNF) adults were all frail except for two individuals who were pre-frail. These older cohorts therefore offered the opportunity to assess the biologic consequences of frailty on platelet function. All participants (including the Young adults) received the seasonal high-dose influenza vaccine, which contains four times the dose of vaccine strain hemagglutinin proteins and is approved for use in adults age 65 and older. We isolated platelet-rich plasma (PRP) from blood samples of participants obtained prior to vaccination (Day 0) and at Day 2, 7, and 28 post-vaccine for isolation of RNA and elucidation of the platelet transcriptome via RNA-seq. Platelet function was further assessed by flow cytometry at Days 0, 2, and 7 post-vaccine in a subset of participants where flow cytometry was done immediately after PRP isolation.

**Table 1:** Cohort characteristics for RNASeq analysis.

| **N = 65** | **Group** |  |  |
| --- | --- | --- | --- |
|  | **Young**<br>**(N = 28)** | **Older (Comm)**<br>**(N = 20)** | **Older (SNF)**<br>**(N = 17)** |
| **Age** |  |  |  |
| Mean (SD) | 28.46 (5.46) | 72.90 (5.47) | 83.53 (11.60) |
| **Biological sex** |  |  |  |
| Female | 18 ( 64.29%) | 11 ( 55.00%) | 9 ( 52.94%) |
| Male | 10 ( 35.71%) | 9 ( 45.00%) | 8 ( 47.06%) |
| **Race** |  |  |  |
| Asian | 5 ( 17.86%) | 0 ( 0.00%) | 0 ( 0.00%) |
| Black or African-American | 3 ( 10.71%) | 0 ( 0.00%) | 0 ( 0.00%) |
| White | 20 ( 71.43%) | 20 (100.00%) | 17 (100.00%) |
| **Hispanic** |  |  |  |
| No | 25 ( 89.29%) | 20 (100.00%) | 17 (100.00%) |
| Yes | 3 ( 10.71%) | 0 ( 0.00%) | 0 ( 0.00%) |
| **Frailty** |  |  |  |
| Non-frail | 28 (100.00%) | 16 ( 80.00%) | 0 ( 0.00%) |
| Pre-frail | 0 ( 0.00%) | 4 ( 20.00%) | 2 ( 11.76%) |
| Frail | 0 ( 0.00%) | 0 ( 0.00%) | 15 ( 88.24%) |
| **Count of conditions\*** |  |  |  |
| Mean (SD) | 0.39 (0.79) | 1.95 (1.82) | 4.41 (2.60) |
| **Medication** |  |  |  |
| Daily Aspirin | 1 ( 3.57%) | 11 ( 55.00%) | 8 ( 47.10%) |
| NSAIDs | 11 ( 39.29%) | 4 ( 20.00%) | 1 ( 5.89%) |
| **Number presc.meds** |  |  |  |
| Mean (SD) | 0.71 (1.21) | 3.40 (2.16) | 9.35 (4.44) |

\*For count of conditions, only conditions occurring in 5% or greater of the sample were included in the count (Congestive heart failure, Coronary Artery Disease, Arrhythmias, High blood pressure, Peripheral vascular disease, Stroke or TIA, Chronic pulmonary disease, Asthma, GERD (Reflux disease), Colitis/Irritable Bowel Disease, Diabetes (requiring medication), Renal insufficiency, Thyroid disease, Anemia).

### Platelets in older adults are characterized by pre-vaccination increased platelet activation and decreased translation pathways

We were interested in associations between pre-vaccination transcriptional profiles and demographic features. The highest proportion of variability in the pre-vaccination state (∼35%) was explained by the group categorization (Young, Older (Comm), or Older (SNF)), with smaller contributions from sequencing run (batch) (∼3%) and biological sex (∼0.7%) (Fig. 1a, Extended Data Fig. 1). However, when considering each group separately, we observed heterogeneity in covariate contribution to the variation, and regressed these covariates in the individual group comparisons (Extended Data Fig. 1). Overall, pre-vaccination RNA expression variability was most highly associated with group membership, and there was a continuum of transcriptional states moving from Young to Older (Comm) to Older (SNF).

**Fig. 1.**
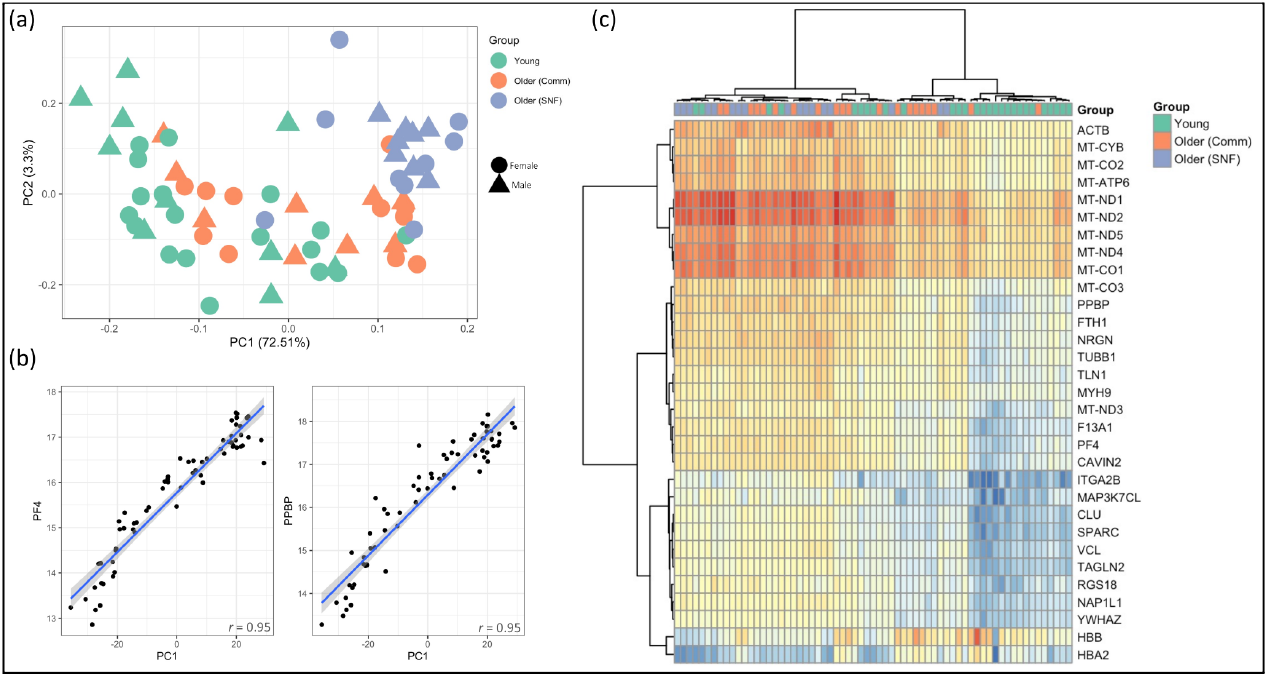
Analysis of RNASeq pre-vaccination data, (a) PCA, (b) correlation of PC1 with platelet activation marker expression, (c) expression heatmap of most variable RNAs (RNAs starting with ’MT-’ are mitochondrial genes).

We performed differential expression (DE) analysis to identify differences in the pre-vaccination transcriptome among the three groups. This analysis found 1508 RNAs that were significantly different when comparing Young v. Older (Comm), and 2209 RNAs that were significantly different when comparing Older (Comm) v. Older (SNF) (|log2fc| > 1.5, adj. p value < 0.05, Table 2). Platelet activation pathways were significantly enriched in Older (Comm) compared with Young adults, as were Rho GTPase effector pathways mediating the activation response^22^ and mitochondrial genes (Table 2, Extended Data Fig. 2a). Notably, activation pathways in platelets from Older (SNF) subjects were also significantly increased compared to Older (Comm) subjects. Overall, these results suggest an increase in platelet activation with increasing age and frailty. Indeed, there was a strong correlation of the first principle component (PC1) (which accounted for over 72% of the variance (Fig. 1a)) with both age and platelet activation (as represented by Platelet Factor-4 (*PF4*) and Pro-platelet Basic Protein (*PPBP*, Fig. 1b).

**Table 2:** Reactome pathways enriched in pre-vaccination platelet RNASeq in (a) Young v. Older (Comm) and (b) Older (Comm) v. Older (SNF) individuals.

| (a) Young v. Older (Comm) |  | (b) Older (Comm) v. Older (SNF) |  |
| --- | --- | --- | --- |
| Enriched in Older (Comm) (n=840) |  | Enriched in Older (SNF) (n=127) |  |
| Pathway | adj. p value | Pathway | adj p value |
| Hemostasis | 2.72E-24 | Hemostasis | 1.65E-05 |
| Platelet activation, signaling and aggregation | 2.72E-24 | Platelet activation, signaling and aggregation | 2.44E-05 |
| Platelet degranulation | 1.02E-16 | RHO GTPases Activate ROCKs | 5.08E-05 |
| Response to elevated platelet cytosolic Ca2+ | 3.35E-16 | Platelet degranulation | 7.09E-05 |
| RHO GTPase Effectors | 2.86E-13 | Response to elevated platelet cytosolic Ca2+ | 7.91E-05 |
| Enriched in Young (n=668) |  | Enriched in Older (Comm) (n=2082) |  |
| Pathway | adj p value | Pathway | adj p value |
| Peptide chain elongation | 2.19E-18 | Cap-dependent Translation Initiation | 8.70E-37 |
| Nonsense Mediated Decay (NMD) independent of the Exon Junction Complex (EJC) | 2.19E-18 | GTP hydrolysis and joining of the 60S ribosomal subunit | 8.70E-37 |
| Viral mRNA Translation | 2.19E-18 | Formation of a pool of free 40S subunits | 8.70E-37 |
| Selenocysteine synthesis | 4.82E-18 | Eukaryotic Translation Initiation | 8.70E-37 |
| Eukaryotic Translation Elongation | 4.82E-18 | L13a-mediated translational silencing of Ceruloplasmin expression | 1.79E-36 |

Additionally, we observed that the most highly expressed RNAs across all subjects had a high proportion of mitochondrial genes, and the mitochondrial genes showed a strong correlation with platelet activation RNA expression (Fig. 1c, Extended Data Fig. 2c).

In contrast to the age-associated increase in platelet activation RNAs, RNAs in pathways related to translation decreased with age and frailty. These pathways were enriched in Young compared to Older (Comm) and Older (Comm) compared to Older (SNF) adults (Table 2, Supplementary Tables 1-4). The use of anti-platelet medications such as non-steroidal anti-inflammatory agents (mainly in the Young group) or aspirin (mainly in the older groups) did not appear to substantially influence these findings, as they were preserved on evaluation of the subset of individuals not on any of these medications, despite the loss in statistical power (Supplementary Tables 5-6). Additionally, a comparison of male versus female differences did not result in any significantly differentially expressed RNAs when all age groups were considered. Taken together, these findings indicate an age- and frailty-associated increase in expression of platelet activation RNAs and mitochondrial genes and a decrease in expression of RNAs encoding translation-related proteins.

Influenza vaccine antibody response was measured using a standard hemagglutination inhibition assay (HAI). To correct for the inverse correlation between pre-vaccine HAI titer and fold-increase post-vaccine, we employed maximum residual after baseline adjustment (maxRBA), a metric which models HAI titer fold changes as an exponential function of strain-specific baseline titers and selects the maximum residual across strains^23^. In order to understand how pre-vaccination transcriptional state is associated with post-vaccination antibody response, we performed a DE analysis of high-vs. low-responders in the pre-vaccination state, and identified 12 up-regulated and 7 down-regulated RNAs in high responders; up-regulated RNAs included *THRB, BLZF1, MPIG6B*, and *CCR1* (Supplementary Table 7). The expression of *THRB*, a nuclear receptor for thyroid hormone, is intriguing in view of the reported role of thyroid hormone on immune response, particularly in the innate immune system^24,25^. The association of *MPIG6B* with vaccine response is notable since it acts as a platelet inhibitory receptor that has also been implicated in early megakaryocyte development^26,27^.

Our previous studies found signatures of high and low vaccine response in PBMCs that differed in different groups^20,23^, and it has been previously shown that sex differences in immune response are also age dependent ^28^. Hence, we assessed what RNAs were DE expressed between males and females and high and low responders in each group, and found that RNAs DE in platelets of males vs. females and high vs. low responders differed substantially for each group (Supplementary Tables 8-10, Supplementary Fig. 1). Examples of RNAs expressed at a lower level in males versus females include the *JUN* oncoprotein in Young adults – previously reported to be subject to regulation by testosterone^29^, and the *MPIG6B* RNA encoding an ITIM-containing platelet inhibitory protein in the Older (SNF) group (also upregulated in high responders across all groups as discussed above). Hence, response-and sex-associated differences in pre-vaccination RNA expression were found to differ between the groups.

### Response to vaccination shows age-specific differential response pathways

To identify patterns within the RNA expression changes in platelets post-influenza vaccination we employed a tensor-based decomposition approach called non-negative CP decomposition (NCPD). This approach models the data as a non-negative sum of rank-one tensors, termed components, each one of which corresponds to a temporal expression pattern shared by a subset of subjects^30^ (Fig. 2). Each RNA and subject has a component score which represents their association with the component time-course pattern.

**Fig. 2:**
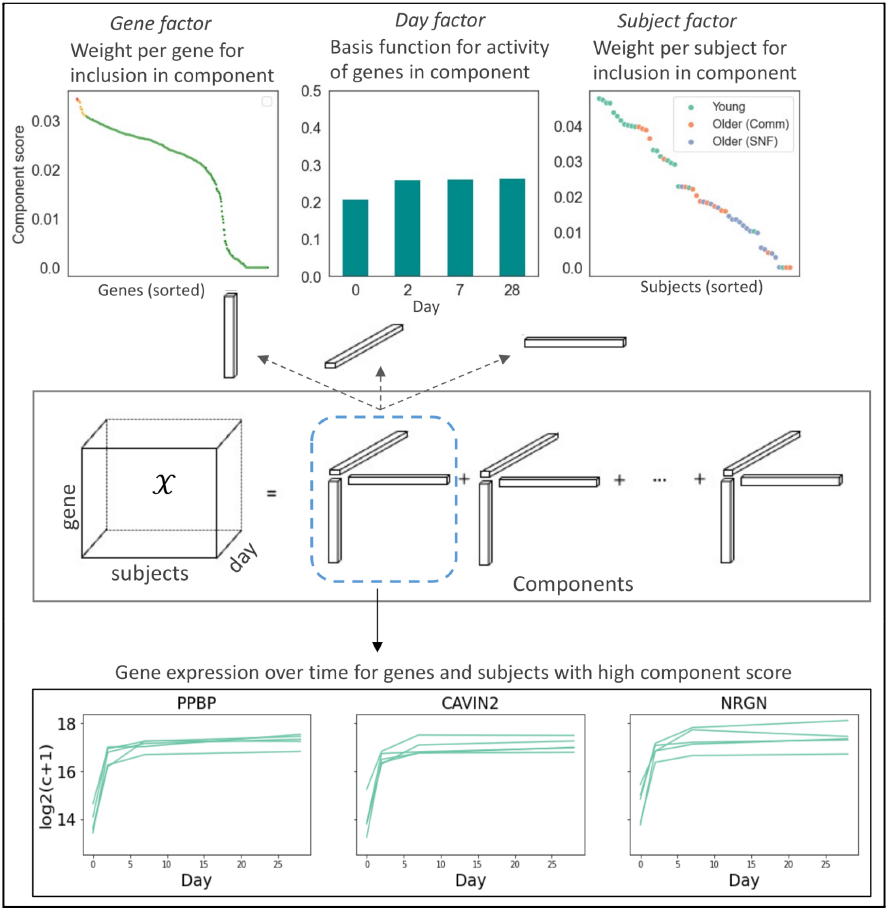
Schematic of non-negative CP tensor decomposition (NCPD) for platelet vaccine response data. The RNA-by-subject-by-day tensor is decomposed into components that represent a time-course pattern of RNA expression for a subset of RNAs across a subset of subjects. Such components can be correlated with activation or deactivation of pathways in specific groups.

The tensor decomposition identified a model with five components. Subject component scores showed a strong association with the three groups (Fig. 3(a,b)), and were significantly correlated with age (Fig. 3c). These associations indicated a strong relationship between transcriptional vaccination response and group and age.

**Fig. 3:**
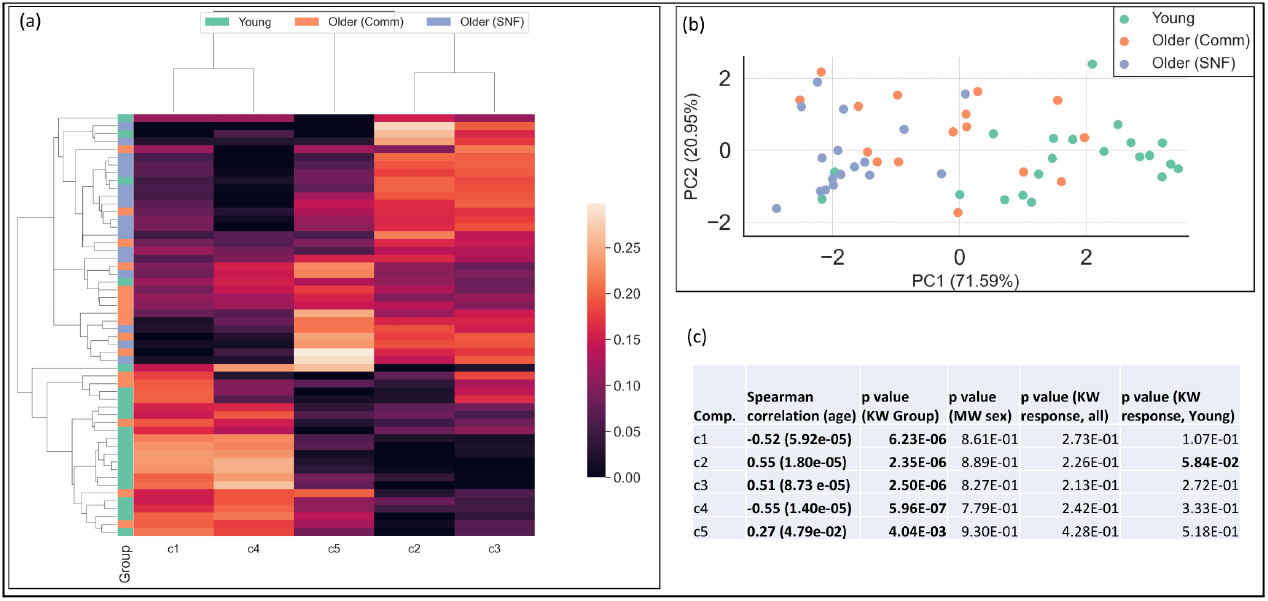
Sample component scores from NCPD of time-course platelet transcriptomic response, (a) Hierarchical clustering and (b) PCA of sample component scores. (c) Association of component scores with age (Spearman correlation) and group (Kruskall-Wallis test), biological sex, and vaccine response (Mann-Whitney U test). Bolded values indicate significance of association (p<0.10).

The top 5% scoring RNAs for Components 1, 2, and 5 were highly overlapping. Pathways involving platelet degranulation, platelet activation, and hemostasis were significantly over-represented. While these components capture similar biology, the temporal patterns and subjects associated with each component differed (Fig. 4). Component 1 was associated with Young adults, and represents a temporal expression pattern where RNA expression increased post-vaccination and remained stable. In Component 2, which was associated with Older (SNF) adults, pre-vaccination RNA levels experienced a sustained drop post-vaccination and in Component 5, pre-vaccination RNA levels experienced a drop followed by a rise back to pre-vaccination levels. The highest scoring subjects in Component 5 were predominantly Older (Comm) adults. The importance of these RNAs in the distinct temporal response between the groups was further supported by differential expression analysis. All 17 of the overlapping RNAs in Components 1,2, and 5 were significantly differentially expressed between at least one group comparison and post-vaccination day in a paired sample analysis (Supplementary Table 12). Overall, these results show that each group displays a distinct temporal pattern of RNAs associated with platelet activation.

**Fig. 4:**
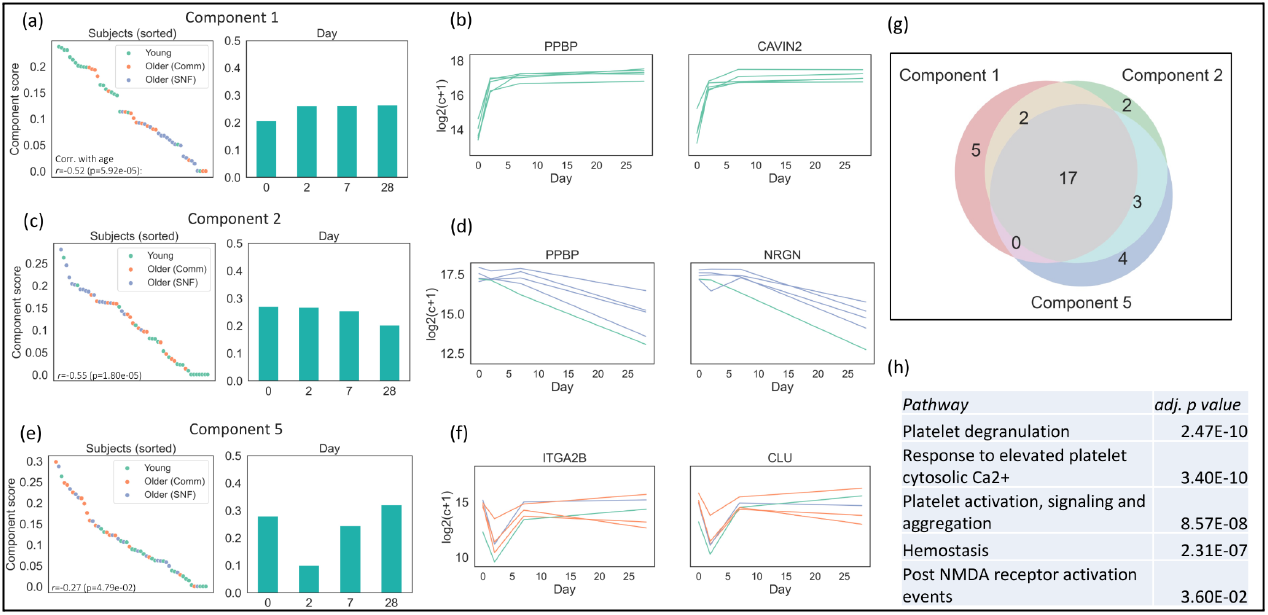
Tensor components related to platelet activation and age group. (a, c, e) Subject and Day scores for Components 1, 2, and 5, respectively, (b, d, f) Expression levels for the top 5 scoring subjects and top 2 scoring RNAs in each component, (g) Venn diagram of overlapping RNAs in Components 1,2, and 5, (h) Over-represented Reactome pathways shared by the three components.

RNAs reflecting expression of mitochondrial genes *MT-1* and *MT-2* were among the top scoring in Components 1 and 2 (Supplementary Table 11), indicating that the correlation between mitochondrial gene and platelet activation RNA expression in the pre-vaccination state may extend post-vaccination. Indeed, the correlation between platelet activation and mitochondrial gene activity was still strongly present in the time-course (Supplementary Fig. 3). Top-scoring RNAs In Components 3 and 4 were also highly overlapping, with pathways involving RNA translation significantly over-represented. We observed that expression of these RNAs tended to have higher pre-vaccination levels which then decreased in Young adults, while pre-vaccination levels were lower and increased in Older (SNF), with intermediate dynamics in Older (Comm) (Extended Data Fig. 3). These RNAs were also found to be significantly differentially expressed in a paired sample analysis (Supplementary Table 12). Therefore, the negative correlation between platelet activation and translation gene RNA levels observed at pre-vaccination were maintained following vaccination.

### Effects of age and frailty on platelet activation at the protein level and comparison with expression of activation marker RNAs following vaccination

We assessed the state of platelet activation at the protein level by determining the expression of p-selectin (CD62p, encoded by the *SELP* gene), CD40 Ligand (CD40L) and CD63 by flow cytometry in Young, Older (Comm) and Older (SNF) groups. All three proteins are found in α-granules (CD62p and CD40L) or dense granules (CD63) within platelets and are transported and expressed on the platelet surface upon activation^31^. We then assessed the expression of RNAs encoding these proteins to evaluate the contribution of transcription to expression of these activation markers.

The levels of surface expression of all three proteins were significantly increased in platelets from Older (Comm), compared to Young adults, suggesting an age-associated increase in platelet activation. We expected that the Older (SNF) group might show similar or even increased expression of these activation markers. However, surface expression in Older (SNF) individuals was significantly decreased compared to Older (Comm), with Older (SNF) activation marker surface expression comparable to that of platelets from Young adults (with the exception of Day 0 CD62p/SELP expression in Older (SNF) residents, which was significantly higher compared to platelets from Young adults) (Fig. 5). The level of these markers in Older (Comm) adults with or without use of anti-platelet medication was similarly increased, indicating that use of anti-platelet medication was not driving this difference.

**Fig. 5.**
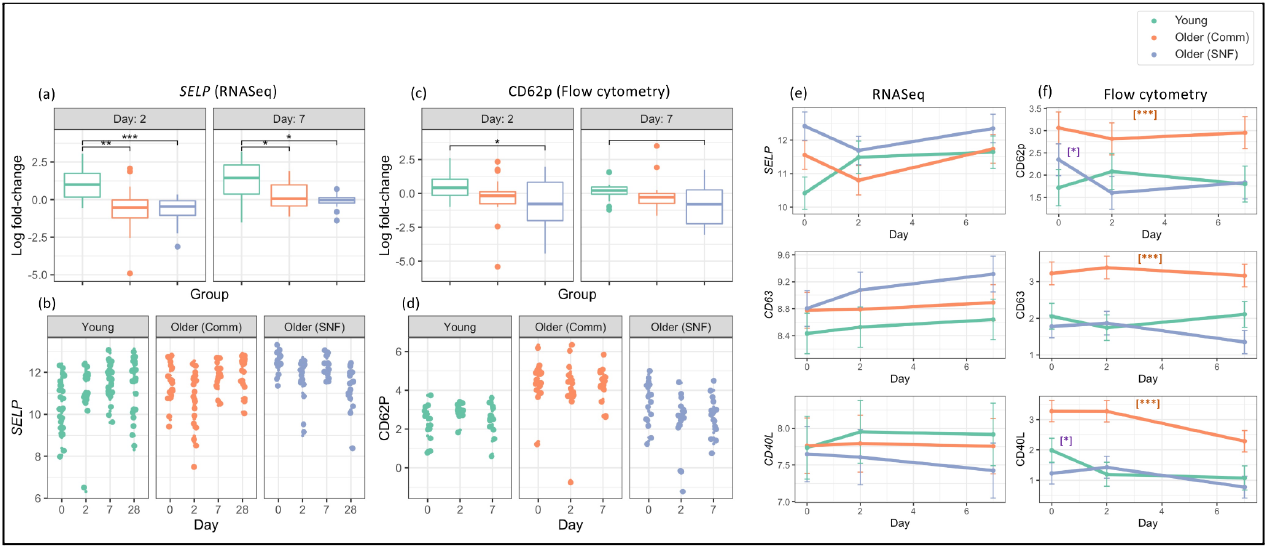
Evaluation of protein and RNA expression of the platelet activation markers p-selectin (CD62p, encoded by the SELP gene), CD63, and CD40L. (a,c) Log2 fold-change at day 2 and 7 compared to pre-vaccination levels of *SELP* (a) or CD62p (c) in Young (RNASeq, n=28; flow cytometry n= 14), Older (Comm) (RNASeq, n=20; flow cytometry, n=17) and Older (SNF) (RNASeq n=17, flow cytometry n=17) adults; (b,d) Scatter plots depicting expression for *SELP* and CD62p; (e,f) Generalized linear mixed effect models for expression of the three markers in (e) RNASeq and of the (f) flow cytometry at pre-vaccination, and days 2 and 7 post-vaccination. Expression coordinates reflect log-transformed normalized expression levels. Significance for (a),(c): ***, p<0.001; **, p<0.01; *, p<0.05; open bracket, p<0.10; significance for (f): orange asterisks indicate the Older (Comm) group was significantly different from Young and Older (SNF) at all days and all time points at adj. p value at least < 0.01; purple asterisks indicate that Young and Older (SNF) were significantly different at day 0 at adj. *p* value at least < 0.05.

We found that expression of the *SELP* RNA at day 2 and day 7 post-vaccine, normalized to day 0, followed the distinct temporal pattern of platelet activation RNA expression identified by the tensor decomposition analysis, with highest fold-changes in Young adults, followed by Older (Comm) and Older (SNF) groups (Fig. 5). The surface expression of CD62p/SELP protein at these post-vaccine time points relative to day 0 closely resembled the RNA expression pattern. We did not observe a similar parallel relationship between normalized RNA expression and protein expression for CD40L and CD63 (Fig. 5). These findings suggest a potential link between RNA and protein expression of *SELP* and CD62p, respectively, despite the presence of pre-formed CD62p within α-granules. Taken together, these findings show an age-associated increase in platelet activation in Older (Comm), compared to Young adults that is attenuated in platelets from Older (SNF) individuals.

### Young individuals exhibit different vaccination dynamics depending on antibody response

We evaluated the relationship between components and influenza vaccine antibody response in all subjects and within each group. The most significant association was found for Component 2, which captured platelet activation among Young adults (p=0.058) (Fig. 6 inset). Among high responders, Component 2 RNAs decreased in expression on days 7-28 whereas in low responders the expression levels remained stable (Figure 6). In order to assess the significance of this observed difference, we performed a DE analysis on the set of top 5% scoring RNAs from Component 2 between Young adult high- and low-responders at each day post-vaccination, and found that more than 50% of the RNAs were significantly DE on day 28, while there were no significant differences at the other timepoints (Fig. 6). In order to observe whether this phenomenon extended beyond the top-scoring Component 2 RNA-set, we performed the DE analysis for all RNAs across the time-course in Young adult high-v. low-responders, and found that the greatest number of differentially expressed RNAs were more highly-expressed in low-responders at day 28. The set of RNAs highly-expressed in low-responders at day 28 encode for proteins involved in pathways that include Rho GTPase effectors and platelet activation (Extended Data Fig. 4). Thus, Component 2 helped to distinguish a difference in time-course patterns of platelet activation RNAs between high- and low-Young adult responders in the day 7-28 post-vaccination period, which extended to additional differentially expressed RNAs involved in Rho GTPase effector pathways in a broader comparison.

**Fig. 6:**
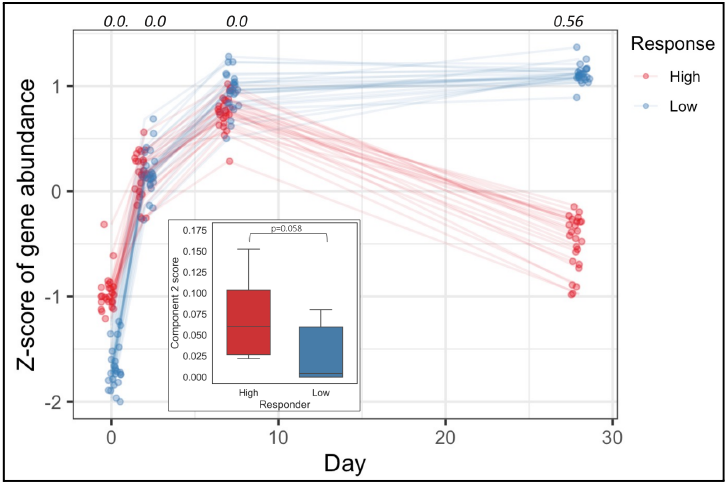
Young adult vaccine high responders show different expression trajectories compared to low responders. Trajectories for high and low-responders of top 5% scoring Component 2 RNAs in young individuals. The fraction of RNAs that are significantly differentially expressed from this set listed on top. Inset: Component 2 score for young high- vs. low responders.

## Discussion

We carried out transcriptomic analyses of human platelets from Young, community-dwelling older (Older (Comm)) and older SNF resident (Older (SNF)) adults in the context of seasonal high-dose influenza vaccination. Using an operational definition of the geriatric syndrome of frailty allowed us to further compare the Older (Comm) group, which was comprised almost entirely of non-frail individuals, to the almost exclusively frail Older (SNF) adults.

Prior to vaccination, we found a marked increase in RNA expression signatures of pathways including platelet signaling, degranulation, and hemostasis in Older (Comm), compared to Young adults and additionally in Older (SNF) residents compared to Older (Comm) adults. These findings indicate that the transcriptome of platelets from non-frail and frail older adults reflects a general activated, pro-thrombotic state compared to young adults. Previous analyses of the platelet transcriptome in healthy young and older, community-dwelling non-frail adults reported evidence for an age-associated increase in Granzyme A expression at the mRNA and protein levels, leading to increased leukocyte signaling and cytokine generation^32^. Our transcriptomic findings indicate an age- and frailty-associated heightened basal activation state in platelets. We also observed increased expression of mitochondrial genes in platelets from both Older (Comm) and Older (SNF), compared to Young adults. Notably, because mammalian platelets are anucleate, the mitochondrial genome represents the only endogenous DNA basally present in platelets. Such mitochondrial DNA encodes 13 proteins mediating oxidative phosphorylation, as well as a group of tRNAs and ribosomal RNAs^33^. Previous studies have revealed increased mitochondrial mass and oxygen consumption associated with TNF-dependent inflammation in aged murine platelets^34^; our findings associated with older and frail older adults suggest a role for mitochondrial dysfunction in human platelets as well. Additionally, our analyses of pre-vaccination platelet RNA expression also revealed a decrease in expression of RNAs related to translation in Older (Comm), compared to Young adults, as well as Older (SNF) versus Older (Comm) adults. Overall, we observed an increase in platelet activation status at the transcriptomic level from Young to Older (Comm) and Older (Comm) to Older (SNF) that was also associated with increased mitochondrial gene expression and decreased levels of RNAs associated with translation. This combination of increased platelet activation and decreased translation in Older (SNF) vs. Older (Comm), and all older compared to Young adults may result in differential functional effects on platelet activation in older adults with or without alterations in functional status.

To analyze temporal patterns we leveraged the TCA tensor decomposition framework that has seen increased application for complex analytics in bioinformatics, including characterizing tissue-specific gene expression phenotypes^35^, *Mycobacterium tuberculosis* subtyping^36^, and systems serology profiling^37^. To extract multi-index patterns that are individually interpretable, we performed a non-negative version of TCA, using CANDECOMP/POLYADIC decomposition (NCPD), which models the data as a non-negative sum of rank-one tensors, termed components^30^. The non-negativity constraint ensures that the model can be analyzed either as a whole and/or on a component-by-component basis since cancellation by negative values will not occur. This is analogous to the strengths exhibited by non-negative matrix factorization (NMF)^38^, which has been used in a variety of biological applications, including sample clustering and biological module detection^39,40^. While other decomposition methods for tensor data exist (Tucker, HGSOC)^41^, NCPD has the advantage that it is both non-orthogonal, which can allow derivation of patterns with overlapping collinear genes, and generates components for straightforward downstream analysis and association with demographic and clinical attributes.

Using a novel application of NCPD, we identified three components corresponding to RNAs that mediate platelet activation, but which showed distinct trajectories among the groups after influenza vaccination – with increased and sustained platelet activation beginning at day 2 in Young adults, decreased activation at day 2 followed by an increase in Older (Comm), and a progressive decrease post-vaccination in Older (SNF) (Fig. 4). Genes associated with these components also included mitochondrial genes, showing that association between platelet activation and mitochondrial RNAs identified at pre-vaccination was maintained following vaccination across groups. We also identified components containing RNAs mediating translation, which demonstrated distinct expression patterns in young versus older groups (Extended Data Fig. 3). Taken together, these results show that platelet activation pathways follow distinct temporal trajectories in the different groups following vaccination, and are associated with broader changes in the transcriptome.

We further studied the effects of age, frailty, and influenza vaccination on proteins associated with platelet activation by assessing the basal expression of surface p-selectin (CD62p, encoded by the *SELP* gene), CD40L, and CD63, prior to and following vaccination using flow cytometry. We found that the change in CD62p protein expression at day 2 and 7 post-vaccine, relative to day 0, closely resembled that observed for normalized *SELP* RNA expression in young, Older (Comm), and Older (SNF) adults at these timepoints (Fig. 5). At both time points post-vaccine, the lowest levels of normalized CD62p surface protein expression were found in Older (SNF) adults, reflecting the generalized decrease in platelet activation following vaccination seen in this group via NCPD (Fig. 4). On the other hand, normalized protein expression of CD40L and CD63 at days 2 and 7 did not parallel expression of their corresponding RNAs. Because CD62p, CD40L, and CD63 surface expression upon platelet activation results from the translocation of preformed protein from alpha granules (CD62p and CD40L) and dense granules and lysosomes (CD63), the parallel trajectories of CD62p surface expression and *SELP* RNA expression suggest a potential contribution of transcription by the platelet parent cell, the megakaryocyte, in the upregulation of surface CD62p protein^42,43^.

In analyzing the absolute levels of CD62p, CD40L and CD63 surface expression pre- and post-vaccine, we found that all three activation markers were markedly elevated in platelets from Older (Comm), compared to Young adults at baseline and at day 2 and 7 post-vaccine – consistent with an age-associated increase in platelet activation (Fig. 5). Interestingly, platelets from the Older (SNF) group showed surface expression levels of these markers that were significantly lower than the Older (Comm) individuals, and were in fact comparable to the Young group (except for a day 0 elevation in CD62p expression that was intermediate between Young and Older (Comm) groups). The basis for this finding remains unclear, and contrasts with the platelet activation RNA expression pattern found in the Older (SNF) group that was markedly increased compared to not only the Young group but also the Older (Comm) group. One possibility is that the substantial decrease in RNAs mediating translation found in the older groups – which was lowest in the Older (SNF) adults – suppressed expression of these activation markers. In this regard, previous studies of the effects of age and frailty on platelet CD62p basal surface expression showed both increased CD62p expression in frail vs. non-frail adults^44^ as well as trends that were similar to our findings^45^. In addition, an age-associated increase in platelet oxidative stress reported in subjects age 40-79 appeared to be reversed in individuals age 80 and above^46^. In sum, the findings of enhanced levels of RNAs associated with activation and mitochondrial gene expression combined with decreased expression of RNAs mediating translation, indicate the presence of alterations in the platelet transcriptome, proteome and activation responses in both non-frail and frail older adults. These changes would be predicted to generally promote a thrombo-inflammatory milieu and may contribute to the increased risk of thrombotic and inflammatory diseases in older and frail older adults.

The basis for age- and frailty-associated alterations in the platelet transcriptome is particularly intriguing since platelets are anucleate and, except for the mitochondrial genome, lack transcriptional activity. As discussed, the possibility that these age-related differences are platelet intrinsic could suggest differences in RNAs expressed in precursor megakaryocytes that generate platelets. Consistent with this hypothesis, other groups have reported that induced changes in platelet RNA expression can be explained, in part, by increased investment of RNA into developing platelets by megakaryocytes^47,48^. Non-cell intrinsic mechanisms such as regulation of post-transcriptional effects on RNA stability by inflammatory stimuli could also play a role^49^; in this context, our finding that influenza vaccine response in Young adults was associated with decreased expression of a component enriched for platelet activation RNAs at day 28 post-vaccine would be consistent with this idea (Fig. 6), since typical human platelet lifespan is approximately 8-10 days^50,51^. Further studies to address this question would provide insights that could be highly relevant to ameliorating not only the substantial burden of cardiovascular and other thrombotic diseases, but also the increased thrombotic complications associated with COVID-19 in older adults.

## Methods

### Human subjects

This study was conducted in accordance with guidelines approved by the Human Investigations Committees of Yale School of Medicine with written informed consent approved annually. Study participants had no acute illness and took no antibiotics or non-steroidal anti-inflammatory drugs within one month of enrollment. Demographic characteristics of participants were collected at enrollment (Table 1). Self-reported information included demographic data, height, weight, medications, and comorbid conditions; immunocompromised individuals were excluded as described previously^52^. Participants were enrolled during the 2018-2019 influenza vaccine season. All participants received the high-dose trivalent influenza vaccine used in that season (Fluzone High-Dose) containing hemagglutinin (HA) proteins from A/Michigan/45/2015 X-275 (H1N1), A/Singapore/INFIMH-16-0019/2016 IVR-186 (H3N2), and B/Maryland/15/2016 BX-69A (a B/Colorado/6/2017-like virus, B Victoria lineage) at a dose of 60µg for each HA. Blood samples were collected prior to vaccination (day 0) and follow-up days 2, 7, and 28. Antibody response to vaccination was assessed using serum samples obtained at day 0 and 28 using a standard hemagglutination inhibition assay^53^.

### Platelet preparation

About 8 ml of blood was collected in acid citrate dextrose (ACD) tubes (Cat. Number 364606, BD Biosciences) for platelet isolation. To avoid shear forces impacting platelet activation, the ACD tube was not drawn first during blood collection. Samples were kept at room temperature and centrifuged at 240xg using a bench top centrifuge (Thermo Fisher Scientific) for 20 minutes without brake. The straw-colored platelet rich plasma (PRP) was carefully transferred to a 15 ml conical tube for RNA extraction and flow cytometry.

### Flow Cytometry

An antibody cocktail for platelet flow cytometry analyses included CD61 FITC (Cat. Number 336404), CD40L PE (Cat. Number 310806, Bio legend), CD14 PE-CF594 (Cat. Number 562335, BD Biosciences), CD63 PercpCy5.5 (Cat. Number 353022, Bio legend), CD41 Alexa Fluor 700 (Cat. Number 303728, Bio legend), CD62p PECy7 (Cat. Number 304922, Bio legend), CD45 APCCy7 (Cat. Number 3368516, Bio legend) and CD66b Pacific Blue (Cat. Number 561649, BD Biosciences). About 100 µl of PRP was mixed with 100 µl of antibody cocktail. After incubation for 20 minutes at room temperature, samples were washed with 1x FACS buffer (1x PBS containing 2% FBS) followed by a paraformaldehyde (PFA) fixation step involving BD Cytofix buffer for 10 min at room temperature. Samples are washed with 1X FACS buffer again to remove the PFA and finally re-suspended in 1 x FACS buffer for flow cytometry analysis using either Fortessa instrument (Becton Dickinson) or CytoFlex LX instrument (Beckman Coulter) fitted with an automated sampler accommodating 96-well plates. FCS files generated by the BD FACS DeVa software (Bd Bio Sciences) or CytExpert software (Beckman Coulter) were analyzed using FlowJo software V10. (FlowJo, LLC). Particles recorded in log scale forward and side scatter (FCS and SSC) were distinguished as anucleate platelets by the surface expression of CD41+ and CD61+. The activation status of platelets was further estimated as percentage of CD41+ CD40L+, CD41+CD62P+ and CD41+CD63+ particles.Samples with excessive aggregation (indistinguishable CD41+CD61+ population) during sample preparation and subsequent staining were excluded from flow cytometry analysis using FlowJo software V10. (FlowJo, LLC). Raw data (fcs files) for the flow cytometry experiments are available via ImmPort (https://www.immport.org) under accession number SDY1393.

### maxRBA calculation

Changes in serology were quantified and vaccine response groups defined using the maximum residual after baseline adjustment (maxRBA) method^23^. Briefly, this approach fits an exponential model to predict titer fold change using baseline titer values for each vaccine strain separately. A subject’s maxRBA score is the maximum residual across all measured vaccine strains for that individual. MaxRBA scores were discretized using quantile cutoffs: those equal to or below the bottom 40th percentile were classified as low responders and those above the top 40th percentile were classified as high responders. We applied this method to generate subject labels separately in the Young and Older(Comm) + Older (SNF) groups.

### RNA Preparation

About 500 µl PRP was added to 700 µl of QIAsol lysis reagent (cat. 217004, Qiagen) and mixed by pipetting at least 10 times to ensure proper lysis. Lysed PRP samples were immediately frozen at -80^0^ C until further extraction. RNA samples were prepared using the miRNeasy kit (cat. 217004, Qiagen) following the manufacturer’s instructions. Briefly, PRP lysed in QIAzol reagent was incubated for 5 minutes at room temperature. To each sample 140 µl of chloroform was added and mixed vigorously and left at room temperature for about 5 minutes. Subsequently, samples were centrifuged at 4^0^ C at 12,000 x g for 15 minutes. The upper aqueous phase containing RNA was carefully transferred to a 2 ml collection tube (cat. 990381, Qiagen) without touching the interphase and placed in a QIAcube instrument for extraction.RNA extraction was carried out using the recommended protocol (FIW-50-001-J_FW_MB and PLC program version FIW-50-002-G_PLC_MB) available from the QIAcube web portal. RNA samples with RNA Integrity Number (RIN) values above 7.0 were used for RNA expression analysis.

RNA-seq libraries were prepared using the Takara Bio SMARTer Stranded Total RNA-Seq-kit - Pico Input Mammalian per the manufacturer’s instructions. Libraries were sequenced on an Illumina NovaSeq 6000, S4 flowcell, 2×100 paired-end, following the manufacturer’s protocols. Low quality reads and reads with length < 50 bp were removed using Trimmomatic v 0.36. Base-quality was then assessed with FastQC v 0.11.7. FASTQ files were aligned using STAR V 2.7.3, against human reference GRCh38p12. Gene counts were determined using HtSeq-count and gencode.v30.chr_patch_hapl_scaff.annotation.gtf. Data for each sample and subject are available via ImmPort (https://www.immport.org) under accession number SDY1393. Raw and count data has been submitted to the Gene Expression Omnibus Database (https://www.ncbi.nlm.nih.gov/geo/) with accession number GSE178158.

### RNASeq processing

The data was pre-processed to only include protein-coding genes and exclude genes on the X and Y chromosome. Following, genes were filtered for low-expression by (1) removing genes with non-zero counts in less than three samples (which corresponds to approximately 1% of total samples) and (2) filtering out the bottom 10% expressing genes of the remainder.

Approximately 16,8000 genes were kept for analysis following pre-processing.

### DESeq Model-based analyses

DESeq2^54^ was used to normalize gene counts prior to performing PCA and hierarchical clustering on the baseline data, as well as for differential expression analysis. For DESeq2 analysis of pre-vaccination data, a Principal Variance Component Analysis (PVCA) was performed on all the groups, as well as each age group to assess covariates to include in the model^55,56^ (Extended Data Fig. 1). For the anti-platelet medication covariate, NSAID use was used for Young subjects, and daily aspirin or prescription anti-platelet medication (‘dAspirinplus’) for the Older (Comm) and Older (SNF) subjects, since these were the predominantly used anti-platelet medications used in the respective groups (Table 1). Covariates that were present in more than 2 subjects were included in the PVCA and models. Batch refers to sequencing run, and cohort for the Young adults the site from which samples were collected.

Based on the results of the PVCA, the model designs were set as follows:

All subjects: ∼ Group + Biological.Sex + Batch,

Young: ∼Biological.sex + Cohort + NSAIDs + Batch,

Older (Comm): ∼Biological.sex + Frailty + dAspirinplus + Batch,

Older (SNF): ∼Biological.sex + dAspirinplus + Batch,

Genes considered as DE had to have a (i) |logFC|≥1.5, (ii) Benjamini-Hochberg adj. p value < 0.05 (unless otherwise noted), (iii) ≥ 25% of samples compared expressing the gene. The latter requirement was necessary in order to exclude genes that were only expressed in a small number of individuals, generally at low counts, which led to very high logFC differences. The functional enrichment analysis on genes considered DE between selected groups was performed on the Reactome database^57^ using g:Profiler (version e104_eg51_p15_3922dba) and Benjamini-Hochberg false discovery rate correction^58^. To assess whether the higher frequency of daily Aspirin use in Older adults may have impacted the group-specific differences, we ran the model design All subjects: ∼Group + Biological.Sex + Batch on subjects that did not use any anti-platelet medication.

DE analysis models for the time-course were built using a multi-level, repeated measures design. Comparisons were made at each day across age groups, and for young individuals, across high- and low-responders.

### Tensor decomposition

Genes for CP decomposition were filtered more stringently than for DESeq2 analysis, since the former does not provide an additional mechanism to address low-expressed genes. Genes were filtered to remove genes with counts ≤ 100 in ≥ 30 samples (which corresponds to approximately 12% of total samples), and the bottom 10% expressing genes were then filtered from the remainder. After filtering, gene counts were normalized using DESeq2 median of ratios method, and *log*_*2*_*(x+1)* transformed. The 500 most variable genes in the count space were used for the decomposition in a tensor framework of genes-by-subject-by-day. Only subjects with data for all days in the study were used. The final dimensions of the tensor were 500 × 54 × 4. The decomposition was performed using CP-OPT with a non-negative lower bound in the Tensor Toolbox package^59^ in MatlabR2020a, which is a gradient-based optimization method that has been shown to be more accurate than CP-ALS (alternating least squares) and more efficient than CP-NLS (non-linear alternating least squares)^59,60^. In order to determine the optimal rank for follow-on analysis, the decomposition was repeated with random initializations, and the normalized Frobenius error and Similarity score were computed for each decomposition (see Supplementary Methods for additional details). A rank 5 model was chosen as it represented the highest Similarity score before a drop with increasing rank and decreased component integrity. Top scoring genes were considered as the top 5% scoring genes for each component based on assessment of a scree plot of gene-by-component scores for each component (Supplementary Fig. 3). All code related to the transcriptional analysis can be found in https://bitbucket.org/kleinstein/projects/src/master/Konstorum2022/.

### Analysis and modeling of flow cytometry markers along with comparison to RNASeq outputs

Longitudinal analysis of flow cytometry and RNA seq marker activation levels were conducted using generalized linear mixed effect models (PROC GLIMMIX) using a lognormal distribution and identity link function. Activation levels for each marker were modeled as a function of group, day of observation, and a group by day interaction adjusted for biological sex, the use of NSAID medication, and daily aspirin use (which corresponds to the same subjects that were on daily aspirin and/or prescription medications as identified by the dAspirinplus category). A spatial exponential covariance structure was included to account for within-participant correlations across repeated measurements at unequal days between observations. Marginal estimates were computed using LSMEANS. These analyses were generated using SAS/STAT software, Version 15.2. Copyright © 2020 SAS Institute Inc. SAS and all other SAS Institute Inc. product or service names are registered trademarks or trademarks of SAS Institute Inc., Cary, NC, USA.

## Acknowledgments

This work was supported in part by awards from the NIH (U19 AI089992 to SHK, RRM, ACS; K24 AG042489 to ACS). The authors gratefully acknowledge the valuable contributions of Dr. John Hwa, colleagues from the Yale Human Immunology Project Consortium (HIPC) group, Bryan Szewczyk and Emma Sykes from the Yale Center for Genome Analysis, Denise Shepard RN, and Mr. Daniel Chawla and Dr. Hailong Meng of the Kleinstein group.

## Author Contributions

A.C.S., S.H.K., and R.R.M. were involved in conceptualization and supervision of the study. M.T.R. provided essential help with platelet biological experiments. A.N. and S.M. recruited participants and collected clinical data and samples. S.M. was responsible for sample preparation, RNA preparation and flow cytometric analyses of platelet function. A.C.S., S.H.K., R.R.M. Y.Z., A.K., B.V.W., and H.G.A. contributed to methodology for analysis and interpretation of results. A.M. performed alignment of the RNAseq data and S.T. and A.M. were responsible for data curation. T.P.B., R.B.B., and D.G.C. performed and analyzed the HAI assays. A.K., B.V.W. and H.G.A. performed a formal analysis of the data. A.K., S.H.K., and A.C.S. drafted the initial manuscript. All authors contributed to revision and editing of the manuscript.

## Extended Data

**Extended Data Fig. 1:**
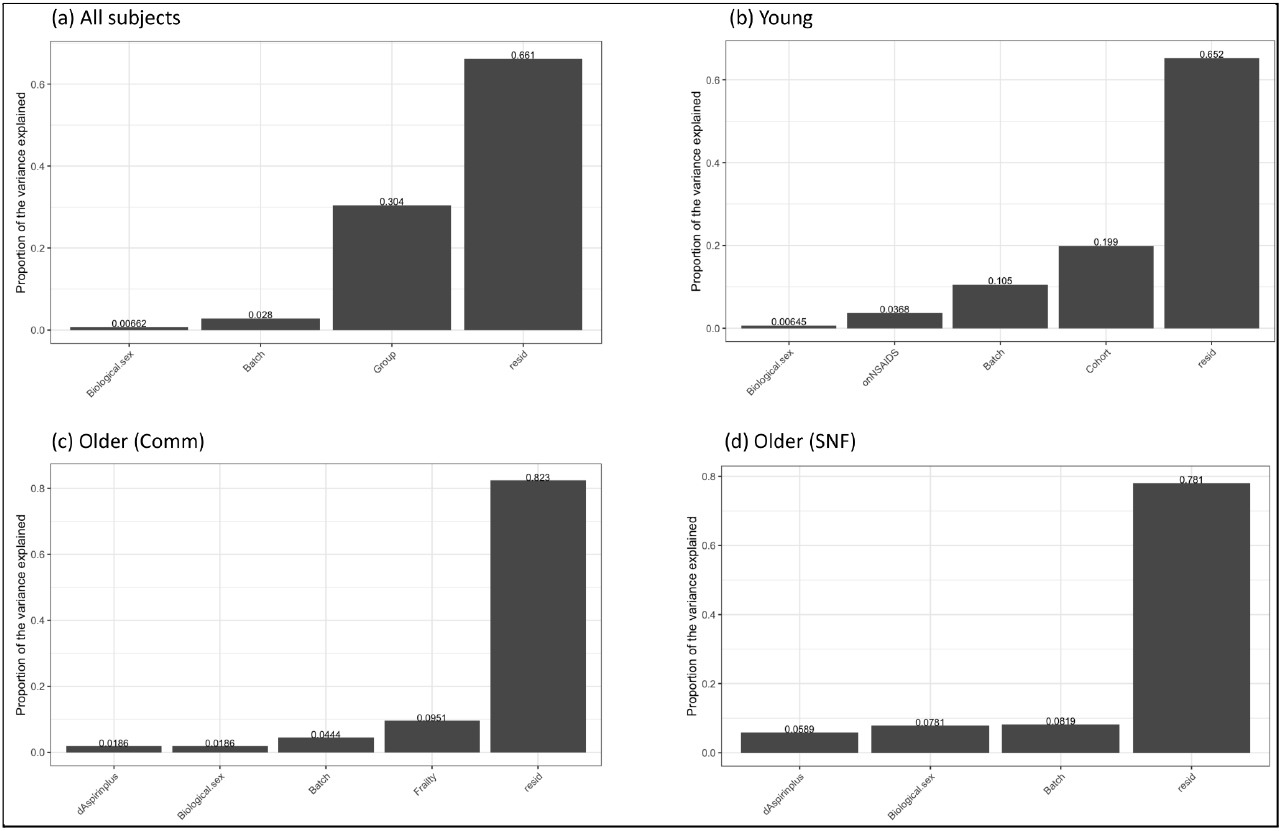
Principal Variance Component Analysis (PVCA) for covariates associated with (a) all subjects and the (b) Young, (c) Older (Comm) and (d) Older (SNF) groups. Biological.sex: biological sex of subject; Batch: sequencing run; Group: Young, Older (Comm), or Older (SNF) group assignment; onNSAIDs: whether subjects was on NSAIDs; dAspirinplus: whether subject was on daily aspirin or prescription anti-platelet medication; Cohort: site location for Young adult sample collection; Frailty: frailty classification (non-or pre-frail) for Older (Comm) adults; resid: undefined residual effects.

**Extended Data Fig. 2:**
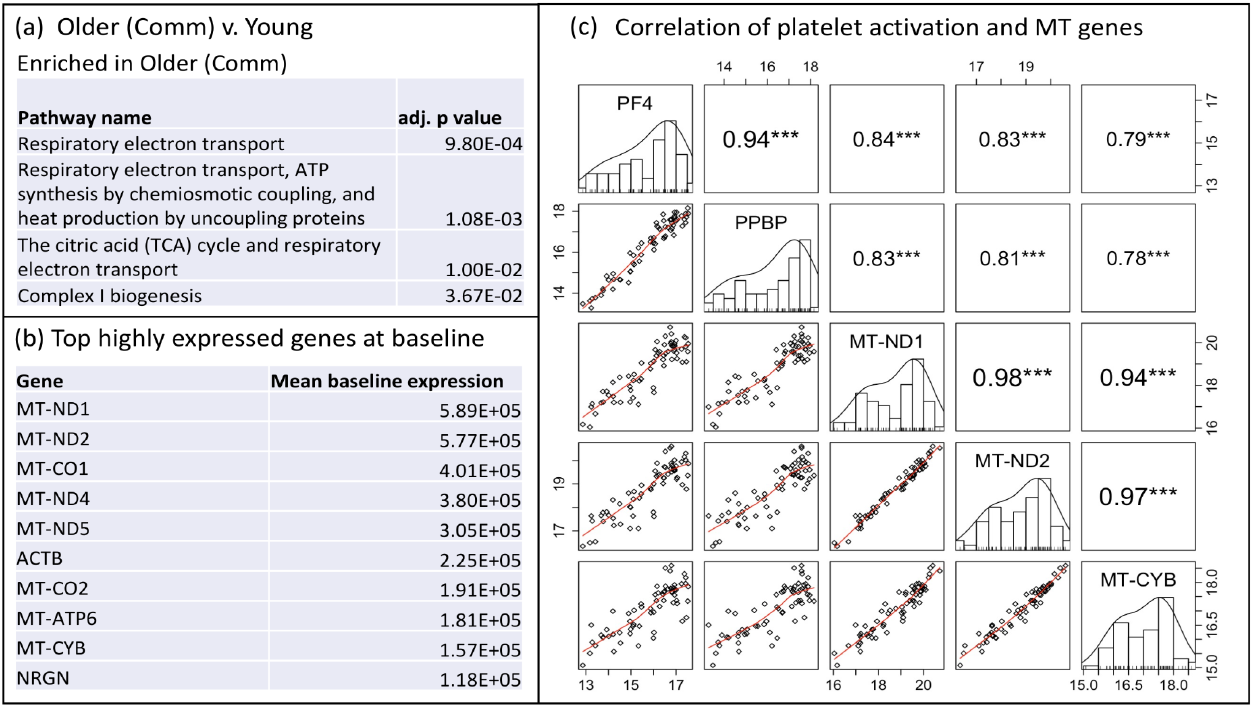
(a) Pathways that involve mitochondrial genes are enriched in Older (Comm) v. Young. (b) Mitochondrial genes are highly expressed in the baseline transcriptome and are (c) correlated with expression of platelet activation RNAs.

**Extended Data Fig. 3:**
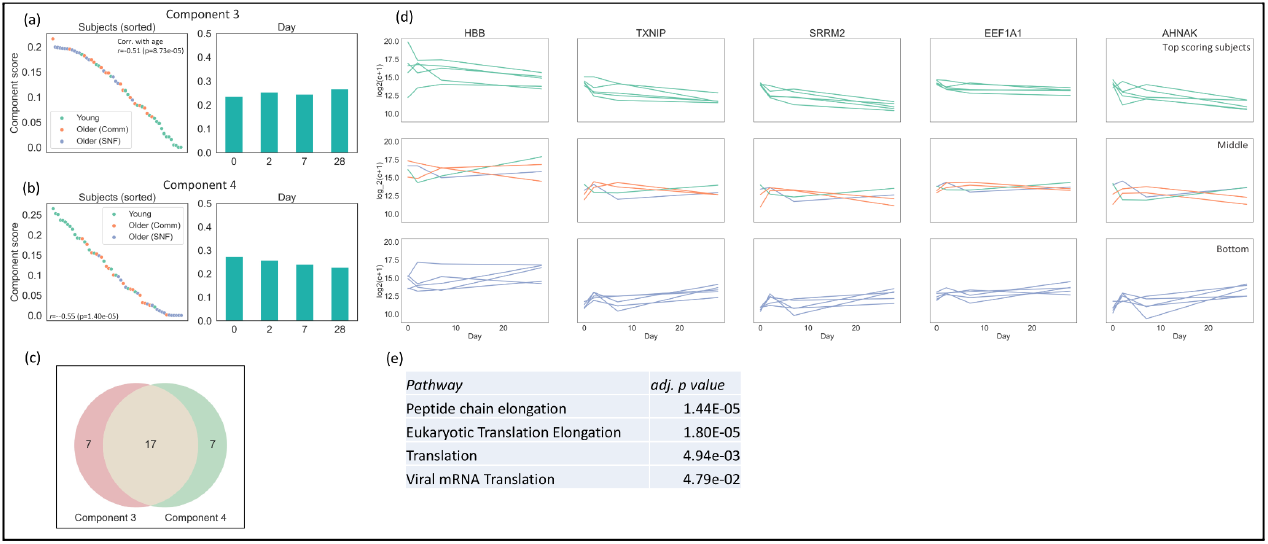
Tensor components relate to translation pathways and age group. (a,b) Sample and Day scores for components 3 and 4, respectively, (c) Venn diagram of overlapping RNAs in Components 3 and 4, (d) Expression levels for the top, middle, and bottom 5 scoring subjects and top 5 scoring RNAs in component 4, (e) Overrepresented Reactome pathways shared by the two Components (adj. p value < 0.05).

**Extended Data Fig. 4:**
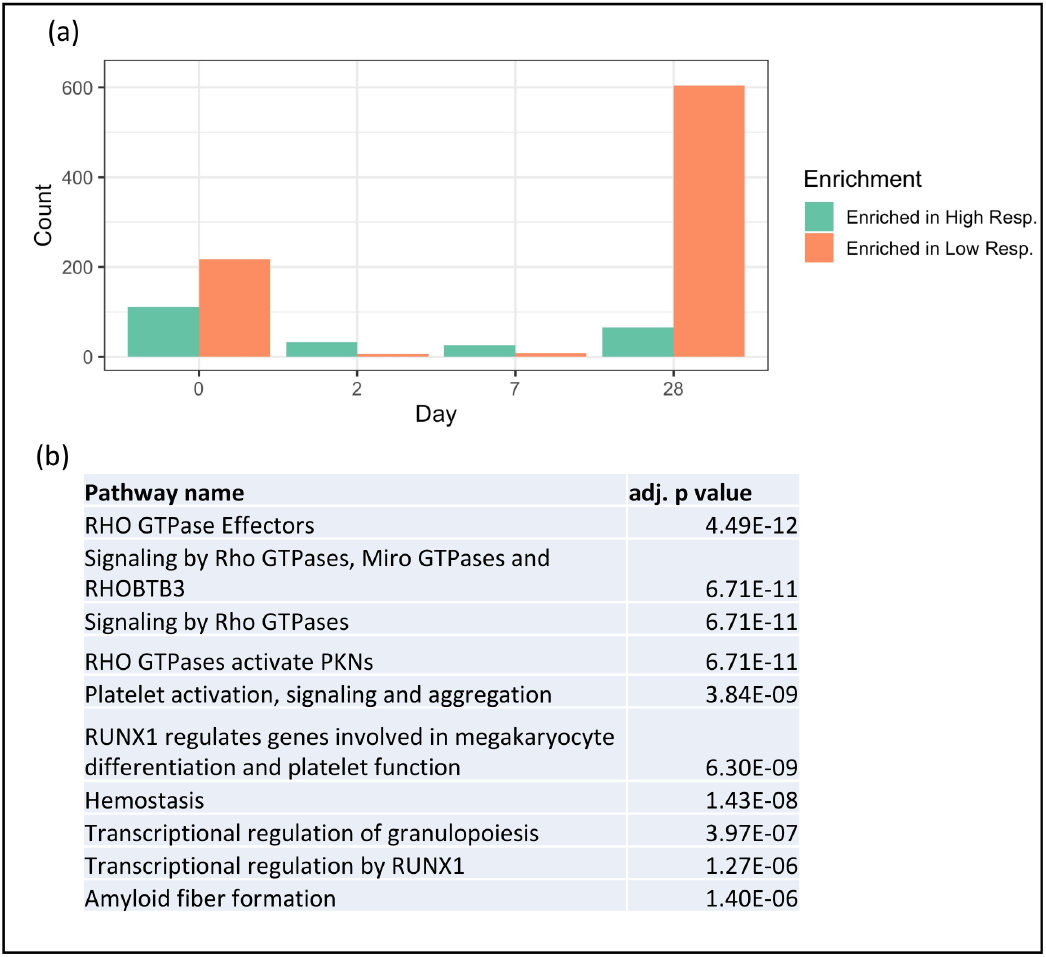
(a) Total number of differentially expressed RNAs (adj. p value < 0.10) between Young high-vs. low-responders for all protein-coding RNAs, (d) Top ten overrepresented pathways of RNAs enriched in low responders at day 28.

## Supplementary Information

**(1) Supplementary_Methods.pdf**

Description of rank and model selection process for Non-negative CP Tensor Decomposition (NCPD)

**(2) Supplementary_Figures.pdf**

Supplementary Figures 1-4.

**(3) Supplementary_Tables.xlsx**

The tabs in the Excel file ‘Supplementary_Tables.xlsx’ correspond to Supplementary Tables 1-11.

**Supplementary Table 1 (ST1_Paths_Enrich_O.CommvYoung)**

Pathways from the Reactome database that are enriched in Older (Comm) v. Young individuals.

**Supplementary Table 2 (ST2_Paths_Enrich_YoungvO.Comm)**

Pathways from the Reactome database that are in Young v. Older (Comm) individuals.

**Supplementary Table 3 (ST3_Paths_Enrich_O.SNFvO.Comm)**

Pathways from the Reactome database that are enriched in Older (SNF) v. Older (Comm) individuals.

**Supplementary Table 4 (ST4_Paths_Enrich_O.CommvO.SNF)**

Pathways from the Reactome database that are enriched in Older (Comm) v. Older (SNF) individuals.

**Supplementary Table 5 (ST5_Paths_Diff_O.Comm_Y(med))**

Pathways from the Reactome database that are enriched Older (Comm) and Young individuals that are not on any anti-platelet medication.

**Supplementary Table 6 (ST6_Paths_Diff_O.SNF_Y(med))**

Pathways from the Reactome database that enriched in Older (SNF) and Young individuals that are not on any anti-platelet medication.

**Supplementary Table 7 (ST7_Genes_DE_Response_All)**

Genes differentially regulated between High- and Low-responders across all age groups.

**Supplementary Table 8 (ST8_Genes_DE_Young)**

Genes differentially regulated in Male v. Female and between High- and Low-responders in Young individuals.

**Supplementary Table 9 (ST9_Genes_DE_O.Comm)**

Genes differentially regulated in Male v. Female and between High- and Low-responders in Older (Comm) individuals.

**Supplementary Table 10 (ST10_Genes_DE_O.SNF)**

Genes differentially regulated in Male v. Female and between High- and Low-responders in Older (SNF) individuals

**Supplementary Table 11 (ST11_NCPD_Component_genes)**

Genes with the top 10% of scores for each NCPD model component. The top 5% genes are used for the downstream analysis.

**Supplementary Table 12 (ST12_Component_genes_DE)**

Significance of DE between top scoring intersecting Component 1,2, and 5 and 3 and 4 genes and all group comparisons at days 2, 7, and 28.

