## Supplementary material for "Platelet response to influenza vaccination reflects effects of aging": Supplementary Figures.pdf

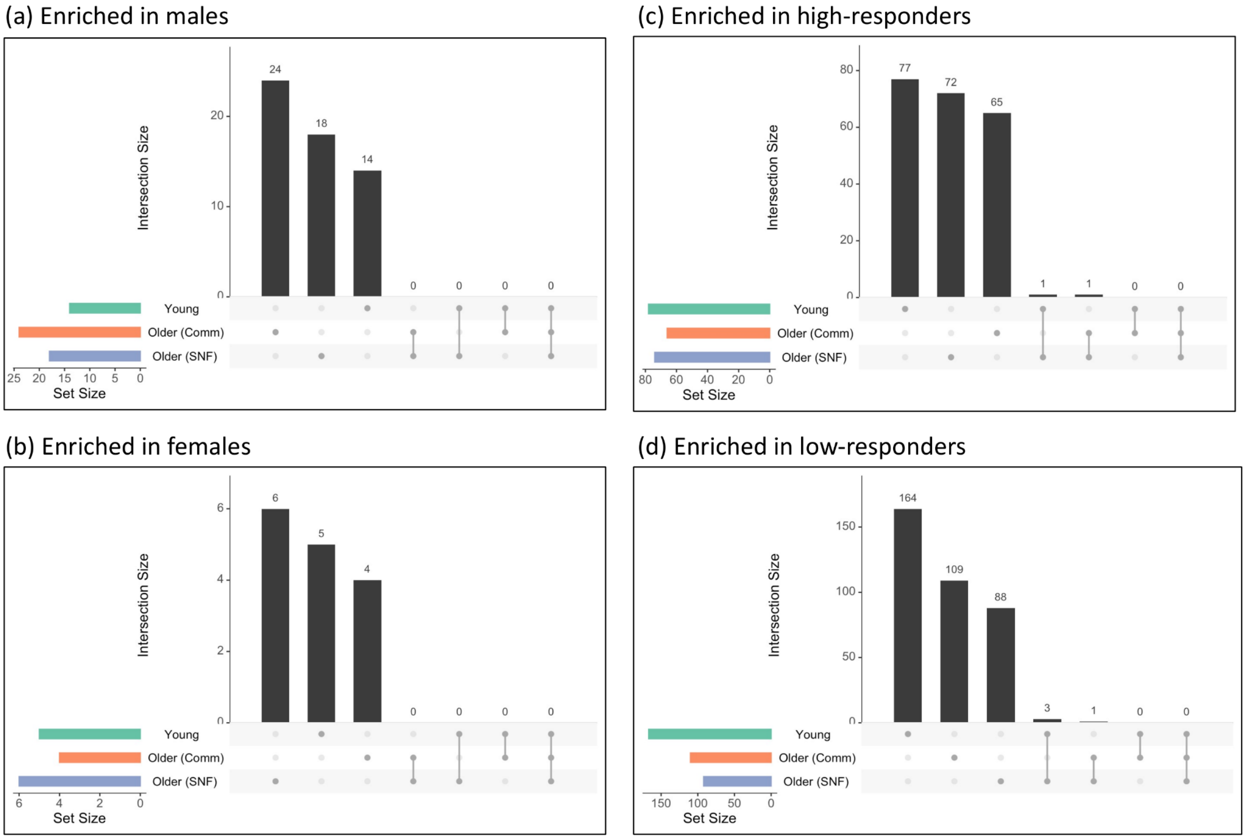


**Supplementary Fig. 1:** Unique and overlapping differentially expressed RNAs at baseline for each of Young, Older (Comm), and Older (SNF) adults with respect to (a) male v. female, enriched in males, (b) male v. female, enriched in females, (c) high- v. low-responders, enriched in high-responders and (d) high- v. low-responders, enriched in low-responders. For (c,d) high- and low-responders counts per group are Young (high, n=7; low, n=9), Older (Comm) (high, n=5; low, n=8 ), Older (SNF) (high, n=5; low, n=4).

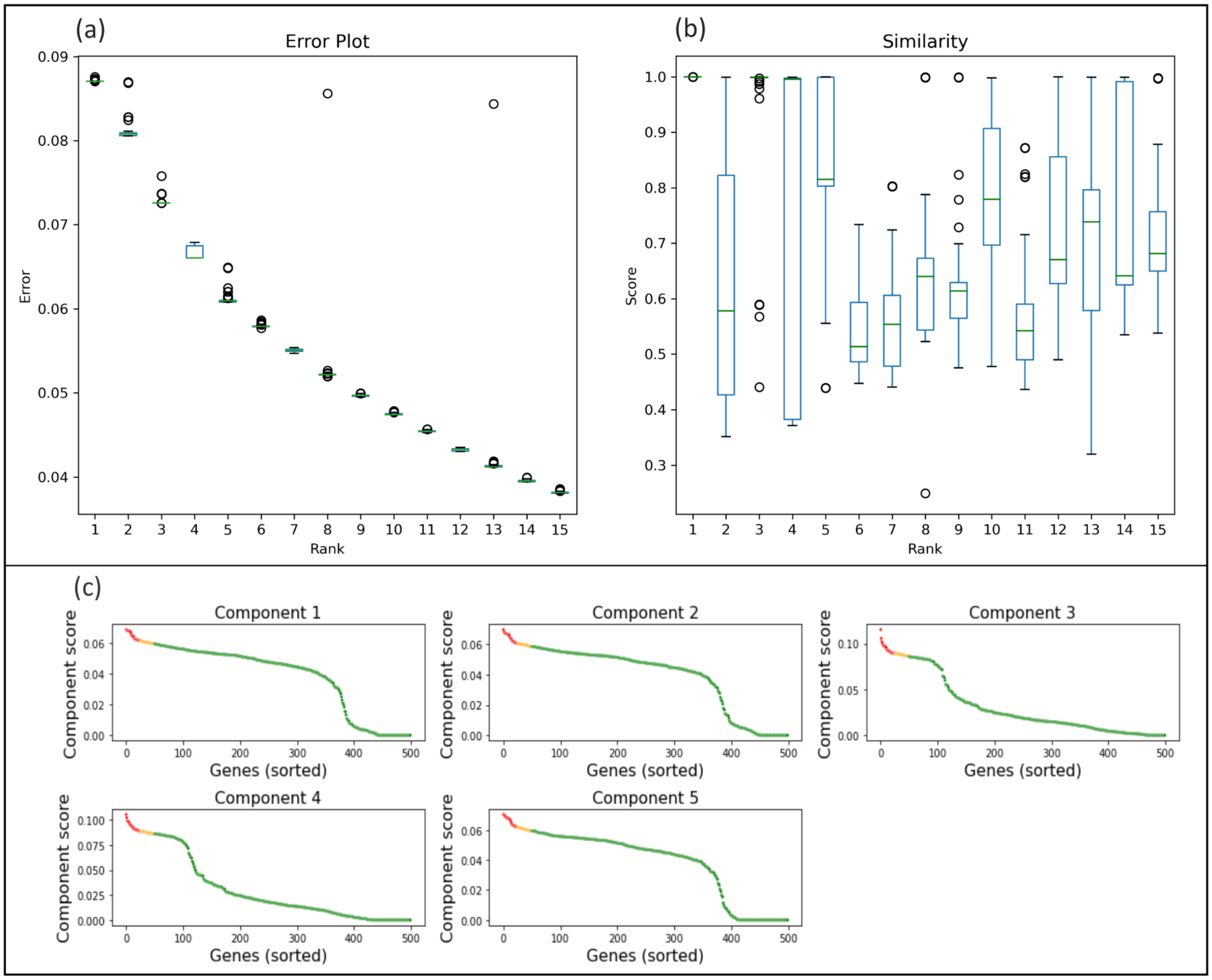


**Supplementary Fig. 2:** Determining optimal rank and model for a time-course platelet vaccination non-negative CP tensor decomposition. (a) Normalized Frobenius error and the (b) Similarity score were calculated for decompositions of ranks $R$=1-15, over 100 random initial conditions for each rank. (c) Gene RNAs ranked by component score for each component of a rank-5 decomposition. Top 5% of scores are in red, top 10% scores are in red and yellow, and remaining scores in green.

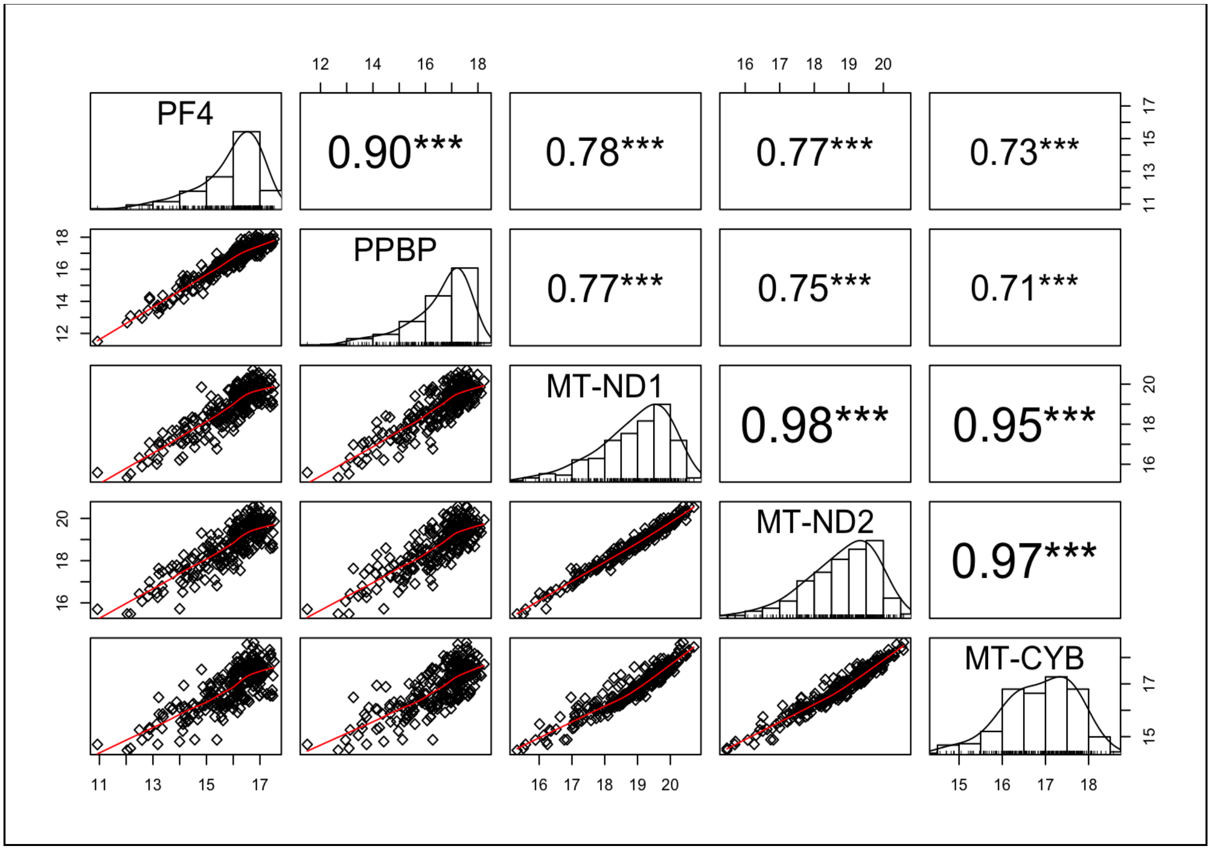


**Supplementary Fig. 3:** Mitochondrial genes, represented here by *MT-ND1*, *MT-ND2*, and *MT-CYB* and platelet activation RNAs, represented here by *PF4* and *PPBP* are significantly correlated across all samples, baseline and post-vaccination.
