## Supplementary material for "Platelet response to influenza vaccination reflects effects of aging": Supplementary_Methods.pdf

### Supplementary Methods
### Platelet response to influenza vaccination reflects effects of aging

Konstorum A., Mohanty S., Zhao Y., Melillo A., Vander Wyk B., Nelson A., Tsang S., Blevins T.P., Belshe R.B., Rondina, M.T., Montgomery R.R., Allore H.G., Kleinstein, S.H., Shaw A.C.

#### Rank and Model selection for Non-negative CP Tensor Decomposition (NCPD)

We performed non-negative CP decomposition (NCPD) in Matlab R2020a using the CP-OPT library in Tensor Toolbox [1], which is a gradient-based optimization method that has been shown to be more accurate than CP-ALS (alternating least squares) and more efficient than CP-NLS (non-linear alternating least squares) [2].

We chose rank based on the the normalized Frobenius error and the Similarity score across 100 random start conditions and decompositions for ranks $R = 1, 2, ..., 15$.

The normalized Frobenius error for a tensor decomposition, $\hat{\mathcal{X}}$, of data tensor, $\mathcal{X}$, can be calculated as

$$\frac{||\mathcal{X} - \hat{\mathcal{X}}||_F^2}{||\mathcal{X}||_F^2}, \tag{1}$$

where $||\cdot||_F$ corresponds to the square root of the sum of squares of elements of a tensor, and is analogous to the Frobenius matrix norm [3].

The Similarity score [4] quantifies the similarity between two models of the same rank by considering the mean of the weighted product of the cosines of matching pairs of loading vectors, where a match is considered as the maximum score across the set all permutations, $\Omega$. For two mode-3 models of rank $R$, $\hat{\mathcal{X}} = [[\lambda^1; A^{(1)}, A^{(2)}, A^{(3)}]]$ and $\hat{\mathcal{Y}} = [[\lambda^2; B^{(1)}, B^{(2)}, B^{(3)}]]$, the Similarity, $S(\hat{\mathcal{X}}, \hat{\mathcal{Y}})$ is calculated as

$$S(\hat{\mathcal{X}}, \hat{\mathcal{Y}}) = \max_{\omega \in \Omega} \frac{1}{R} \sum W_{r,\omega}(\lambda^1, \lambda^2) \prod_{i=1}^{3} {a_r^{(i)}}^T b_{\omega(r)}^{(i)}, \tag{2}$$

where $W_{r,\omega}(\lambda^1, \lambda^2) = (1 - |\lambda_r^1, \lambda_{\omega(r)}^2| / \max(\lambda_r^1, \lambda_{\omega(r)}^2))$ penalizes a difference in weights between two components, especially when the weights are of large magnitude. Thus, $W$ penalizes two models that may have otherwise similar components but have been weighted differently.

Model selection was based on identifying a rank that produces the smallest normalized Frobenius error while maintaining a high Similarity score. Additionally, component integrity was be examined to investigate whether expression patterns corresponded in a one-to-one fashion to component temporal patterns.
